# High-resolution transcriptomic profiling of *Arabidopsis thaliana* across a 37°C–40°C thermal gradient

**DOI:** 10.64898/2026.07.19.739423

**Authors:** Elisabeth Gillham, Jason Hu, Yifan Huang, Aimi Kochi, Kiran McDonald, Caleb Scott-Joseph, Chenhao Xu, Ming Ye, Nick Kaplinsky

**Author notes:** These authors contributed equally to this work (Elisabeth Gillham, Jason Hu, Yifan Huang, Aimi Kochi, Kiran McDonald, Caleb Scott-Joseph, Chenhao Xu, Ming Ye).

## Abstract

Thermal adaptation is critical for organismal viability, with acquired thermotolerance (AT) in *Arabidopsis thaliana* typically conferred by acclimation temperatures between 34°C and 37°C, while acclimation at 40°C fails to protect against lethal heat stress. To elucidate the transcriptional mechanisms underlying the loss of AT at higher temperatures we performed RNA-seq profiling of Arabidopsis seedlings across a single-degree thermal gradient from 37°C to 40°C. Our analysis reveals that all of these temperatures result in a robust heat shock response, characterized by the upregulation of genes involved in protein folding and stress responses. However, each temperature elicits a distinct transcriptional signature. These findings demonstrate that temperature-specific fine-tuning of the heat shock response occurs within this narrow range and that these differences may dictate the successful acquisition of thermotolerance. This dataset provides a valuable resource for understanding the molecular architecture of heat-stress adaptation and the distinct transcriptional states associated with different thermal regimes.

## Background & summary

Thermal adaptation modulates developmental trajectories, organismal viability, and agricultural productivity in crop species (Chen et al. 2022). The molecular architecture governing the adaptation pathways that confer thermotolerance are well-established and include regulatory networks of transcription factors including heat shock factors (HSFs) and induced proteins that mediate cellular stress responses, predominantly heat shock proteins (HSPs). The conserved transcriptional heat shock response (HSR) is activated in response to thermal conditions that induce proteotoxic stress, including protein denaturation and aggregation (Li et al. 2025). The HSR is mechanistically distinct from the temperature compensation machinery utilized during moderate but non-stressful temperatures (Zhou et al. 2022).

Exposure to a one-hour 45°C heat shock (HS) is lethal to non-acclimated Arabidopsis Col-0 seedlings. Acclimation at 37°C before an otherwise lethal HS provides robust thermotolerance, a phenomenon called acquired thermotolerance (AT) (Silva-Correia et al. 2014).

High-temperature regimes with different magnitudes and timing elicit fine-tuned transcriptional and organismal responses (Yeh et al. 2012). An example of this regulatory fine-tuning of the HSR is the differential transcriptional and physiological behavior observed between gradual and step-wise temperature increases. Although both treatments result in upregulation of a shared set of HSR genes, gradual temperature increases result in significant transcriptional differences accompanied by higher organismal survival compared to step-wise increases (Larkindale and Vierling 2008). A second example shows that small temperature differences can have large effects on organismal thermotolerance. In Arabidopsis, thermotolerance can be acquired with acclimation temperatures between 34°C–37°C while acclimation at 40°C, a temperature that is not inherently lethal, fails to generate AT (Silva-Correia et al. 2014).

To investigate HSR fine-tuning across the temperature range where AT is lost, we used RNA-seq to generate Arabidopsis transcriptional profiles across a thermal gradient spanning 37°C to 40°C with single degree resolution. Our analysis indicates that although each of these temperatures elicits a robust HSR, temperature-specific differences in gene expression clearly distinguish each temperature. Understanding these differences may provide insights into specific HSR patterns that dictate the successful acquisition of thermotolerance.

## Methods

### Plant growth and RNA extraction

For each sample, 100mg of Col-0 seeds were surface sterilized with an ethanol rinse followed by 10’ in a 30% bleach, 0.1% SDS solution and then five rinses in sterile water. Sterilized seeds were stratified for two days at 4°C and then transferred to 50ml of 0.5x MS in a 250ml flask and grown under constant light (150 μmol m^−2^ s^−1^) at 20°C on a 100rpm shaker. After five days of growth, three flasks of seedlings per treatment temperature were transferred to recirculating water baths (A2.2, Anova Applied Electronics, Inc.) set to 22°C, 37°C, 38°C, 39°C, or 40°C. Water temperatures were monitored using a calibrated RTD sensor thermometer (THS-222-555, ThermoWorks Inc.) and did not deviate from the set temperature by more than 0.05°C. After a one hour treatment, 50mg of plant material was flash frozen in liquid nitrogen and used for RNA extraction. Total RNA was extracted using a RNeasy Plant Mini Kit and treated with an on-column RNase-Free DNase Set (Qiagen N.V.).

### RNAseq library preparation and sequencing

RNA sample quality was assessed by High Sensitivity RNA Tapestation (Agilent Technologies Inc.) and quantified by Qubit 2.0 RNA HS assay (ThermoFisher). Samples had RIN values between 7.5 and 8.2. Sequencing libraries were prepared using NEBNext Poly(A) mRNA Magnetic Isolation Module and NEBNext Ultra II Directional RNA Library Prep Kit for Illumina (New England BioLabs Inc.). Libraries were sequenced on an Illumina NovaSeq 6000 (Illumina Inc.) with a read length configuration of 150 PE for 40M reads per sample (20M in each direction).

### Sequence analysis

Fastq files were trimmed and filtered for low-quality sequences using Trimmomatic v0.38 [ILLUMINACLIP:NexteraPE-PE.fa:2:30:10:8:true SLIDINGWINDOW:4:25 HEADCROP:15] (Bolger et al. 2014). Trimmed and filtered Fastq files were then mapped to the TAIR10 reference genome using HISAT2 v2.2.1 [--trim5 ‘0’ --trim3 ‘0’ --phred33 --pen-cansplice 0 --pen-noncansplice 12 --pen-canintronlen G,-8.0,1.0 --pen-noncanintronlen G,-8.0,1.0 --min-intronlen 20 --max-intronlen 5000] (Kim et al. 2015). Araport 11 annotations (Cheng et al. 2017) were used to generate gene count files with FeatureCounts v2.0.3 (Liao et al. 2014). Differentially expressed genes were identified using DESeq2 v1.40.2 (Love et al. 2014). Functional enrichment analyses were performed using gProfiler (Raudvere et al. 2019).

### Data record

Raw fastq files have been deposited at NCBI SRA as BioProject ID PRJNA1481274 containing BioSamples SAMN61073994-SAMN61074008 and SRA records SRR39287100-SRR39287114. BioSamples and SRA files are named with an XXY convention where XX is the temperature (22°C, 37°C, 38°C, 39°C, or 40°C) and Y is the replicate (A, B, or C).

## Technical Validation

### Sequencing quality metrics

After trimming and quality filtering, each sample contained an average of 30.6M paired-end reads (27.3M-33.4M) with an average Phred quality score of 34.9 (Fig 1). An average of 85.6% (83.1%-87.2%) of read pairs mapped concordantly exactly once representing between 23.6M-28.1M paired-end reads per sample.

**Figure 1.**
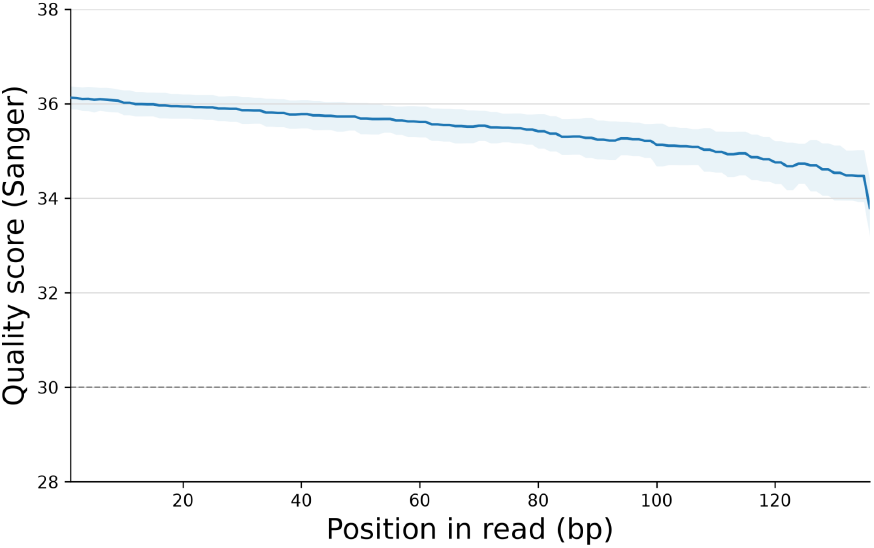
Mean Sanger quality score across all 15 samples by read position; shading indicates ±1 SD and the dashed line indicates Q30.

### Principal Component Analysis (PCA) and Sample Distance Analysis

To evaluate the reproducibility of our biological replicates and identify global differences between treatment groups, we performed Principal Component Analysis (PCA) and hierarchical clustering based on pairwise Euclidean distances. PCA revealed clustering of replicates within experimental groups and robust separation between experimental groups primarily along the first principal component (PC1), which accounted for 88% of the total dataset variance (Fig. 2A). This primary axis cleanly separates the 22°C samples from the 37°C-40°C samples and also clearly distinguishes the 37°C-40°C samples from each other. This demonstrates that temperature was the dominant driver of transcriptional variation and that single degree temperature differences result in distinct transcriptional states. Clustering by sample distance also shows that replicates cluster with each other and that each temperature results in a distinct transcriptional state. Among the high-temperature treatments, 40°C is globally most similar to 22°C, followed by 39°C and 38°C. While 37°C is the closest temperature to 22°C, the 37°C and 22°C transcriptomes are less similar to each other than any other pairwise comparison (Fig. 2B).

**Figure 2.**
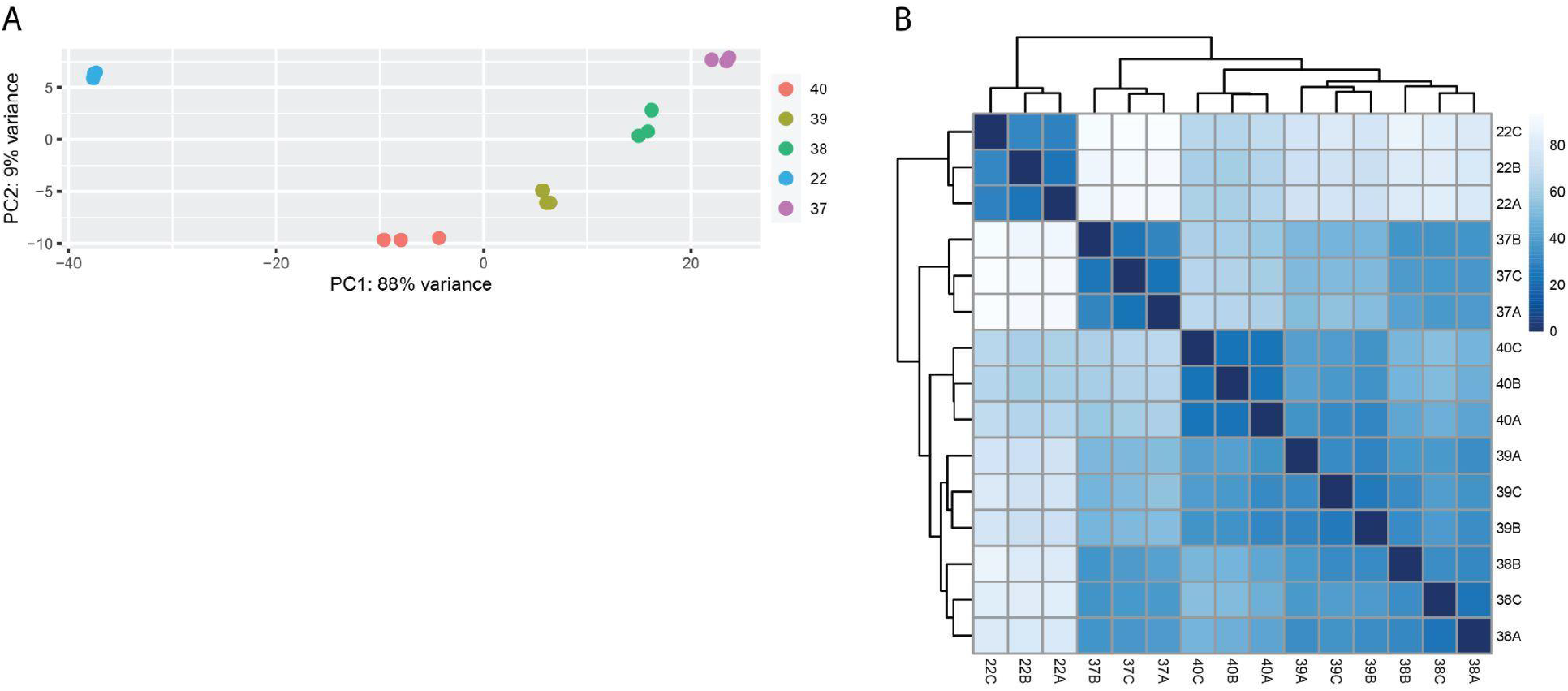
Principal component analysis of normalized gene expression (A). Hierarchically clustered sample-to-sample distance matrix (B).

### Functional Enrichment Analysis

To determine if all four high-temperature treatments resulted in a HSR, we identified the genes induced at each elevated temperature vs 22°C (p_adj_<0.001, log_2_ FC>1) and performed a Gene Ontology (GO) enrichment analysis on them. This analysis revealed remarkable uniformity across the entire thermal gradient. Each high temperature resulted in induction of a set of identical high-level HSR-relevant Biological Process (BP) and Molecular Function (MF) terms (Table 1). Key enriched terms included response to heat (GO:0009408), unfolded protein binding (GO:0051082), protein folding chaperone (GO:0044183), and heat acclimation (GO:0010286). These results demonstrate that the HSR in Arabidopsis is fine tuned in response to small differences in temperature, that this tuning is associated with important biological outcomes (survival vs death), and finally that insights into this functional tuning may not reliably emerge from standard functional enrichment approaches.

**Table 1.** Enrichment of HSR related molecular function and biological process GO terms. Adjusted p-values are shown for enrichment of genes expressed at elevated temperatures relative to 22°C.

| GO term | 37 vs 22 | 38 vs 22 | 39 vs 22 | 40 vs 22 |
| --- | --- | --- | --- | --- |
| MF: Protein folding chaperone | 2.1x10<sup>-22</sup> | 7.7x10<sup>-23</sup> | 3.2x10<sup>-19</sup> | 1.6x10<sup>-11</sup> |
| MF: ATP-dependent protein folding chaperone | 3.8x10<sup>-22</sup> | 2.2x10<sup>-22</sup> | 6.1x10<sup>-21</sup> | 6.7x10<sup>-14</sup> |
| MF: Unfolded protein binding | 3.8x10<sup>-10</sup> | 8.7x10<sup>-10</sup> | 1.9x10<sup>-8</sup> | 1.1x10<sup>-6</sup> |
| MF: Protein-folding chaperone binding | 4.3x10<sup>-10</sup> | 1.6x10<sup>-10</sup> | 9.4x10<sup>-11</sup> | 9.4x10<sup>-11</sup> |
| MF: Heat shock protein binding | 2.0x10<sup>-5</sup> | 5.0x10<sup>-5</sup> | 7.6x10<sup>-5</sup> | 6.6x10<sup>-4</sup> |
| MF: Hsp70 protein binding | 1.3x10<sup>-3</sup> | 2.6x10<sup>-3</sup> | 3.1x10<sup>-3</sup> | 4.2x10<sup>-3</sup> |
| BP: Response to heat | 3.0x10<sup>-61</sup> | 2.8x10<sup>-61</sup> | 8.6x10<sup>-63</sup> | 7.3x10<sup>-53</sup> |
| BP: Response to stress | 8.5x10<sup>-36</sup> | 1.3x10<sup>-32</sup> | 1.2x10<sup>-33</sup> | 2.9x10<sup>-28</sup> |
| BP: Protein folding | 1.3x10<sup>-29</sup> | 1.2x10<sup>-27</sup> | 6.0x10<sup>-25</sup> | 1.7x10<sup>-18</sup> |
| BP: Cellular response to stress | 2.9x10<sup>-28</sup> | 5.0x10<sup>-27</sup> | 2.4x10<sup>-27</sup> | 3.5x10<sup>-26</sup> |
| BP: Heat acclimation | 3.6x10<sup>-22</sup> | 2.1x10<sup>-22</sup> | 3.1x10<sup>-21</sup> | 5.2x10<sup>-19</sup> |
| BP: Cellular response to heat | 4.6x10<sup>-6</sup> | 1.9x10<sup>-6</sup> | 2.6x10<sup>-6</sup> | 1.3x10<sup>-6</sup> |

